# Quantification of cardiac-locked brainstem velocity at high resolution based on retrospectively-gated DENSE MRI at 7T

**DOI:** 10.64898/2026.07.06.736820

**Authors:** Amelia Strom, Zijing Dong, Timothy G. Reese, Laura D. Lewis, Jonathan R. Polimeni

## Abstract

**Purpose:** The motion of the brain tissue within the skull is thought to be induced by cardiac pulsations and other hemodynamic processes and may influence CSF flow, but its precise drivers and downstream effects are unclear. Understanding these phenomena requires an accurate and precise method of tissue motion quantification that can be extended to investigate multiple potential drivers of tissue motion *in vivo*.

**Methods:** Here, a version of the Displacement ENcoding with Stimulated Echoes (DENSE) pulse sequence was implemented to measure cardiac-locked velocity responses in the pons and midbrain in eight healthy volunteers. The method featured retrospective cardiac gating, a single mixing time, and measured multiple voxel sizes.

**Results:** A previously undescribed double-peak pattern of cardiac-locked longitudinally directed brainstem velocity was identified that appears to reflect both nonrigid and rigid motion components. This pattern was only visible when estimating the cardiac response using absolute time after systole instead of the percentage of the cardiac cycle. Measurements were performed in a custom-built slow-flow phantom, and repeat sessions were acquired in two volunteers to assess accuracy and precision. Despite potential increased influence of CSF motion with larger voxel sizes, no voxel-size-dependent bias was found in the velocity estimations. Lack of voxel size bias was attributed to the complexities of partial-volume effects between CSF and tissue in phase-valued data that were evaluated using numerical simulations.

**Conclusion:** In sum, a method to measure brain tissue motion with high spatiotemporal precision is presented that can be extended to applications beyond measuring cardiac-locked motion.

## Introduction

The brain parenchyma is a soft, deformable tissue that undergoes displacement as a mechanical response to various physiological dynamics. While brain tissue motion has been observed with a variety of methods^1–4^, its precise drivers and downstream effects are not well understood. Processes such as the cardiac cycle, respiration, neuronal activity, and locomotion generate tissue motion that may be driven by changes in cerebral blood volume or intracranial pressure^2,5,6^. One potential downstream effect of tissue motion is the consequent flow of interstitial fluid or CSF within and around the tissue, which is thought to be essential for brain waste clearance. Several waste products including amyloid-β are thought to be transported by CSF within perivascular spaces towards the subarachnoid space and subsequently out of the cranium^7^. Brain tissue motion may thus be relevant for fluid flow and clearance at multiple spatial scales. At the scale of the perivascular space, computational models must account for the mechanical properties of brain tissue surrounding these fluid-filled spaces to accurately model periarterial or perivenular flow^8^. At the scale of the whole brain, tissue motion has been shown to be coupled to ventricular CSF flow over the cardiac cycle^9^. Both the efficiency of brain waste clearance^10–12^ and the viscoelastic properties of brain tissue degrade with age^13–15^, suggesting that tissue displacement may play a role in age-dependent clearance dysfunction.

Understanding the drivers and consequences of tissue motion requires accurate and precise quantification, especially given the very slow velocity of the tissue. Also, noninvasive measurement is required, since invasive techniques will disrupt intracranial pressure and the mechanical dynamics within the cranium. *In vivo* MRI is well suited for tissue motion quantification; indeed, multiple MRI-based tissue motion estimation methods have been used over decades^16–23^, such as gradient-echo-based phase-contrast^16,24–26^, amplified MRI^19^, and non-balanced SSFP^27^. Here, we evaluated the use of Displacement ENcoding using Stimulated Echoes (DENSE)^28^ due to its ability to quantify subtle tissue motion; the stimulated echoes employed by DENSE uniquely allow for very long motion-encoding times because the magnetization decays with T_1_ over the mixing time. Thus, DENSE can be used to quantify motion dynamics occurring over long timescales to investigate tissue motion drivers other than the cardiac cycle.

DENSE has been used previously to measure brain tissue motion over the cardiac cycle^29–35^ and respiratory cycle^29^, and to quantify tissue strain metrics^9,29,30,33^. Deep gray matter regions like the brainstem are particularly well studied^30–32^, since the increase of cranial blood volume during the systolic phase of the cardiac cycle causes relatively large amplitude displacement of the brainstem through the foramen magnum^37^, consistent with the Monro–Kellie doctrine^38^. However, previous DENSE implementations have used prospective cardiac gating, limiting temporal precision and impeding applications beyond cardiac-locked motion quantification. Prospective gating also causes fluctuations in signal magnitude by dynamically altering the TR and thus the amount of T_1_-related longitudinal relaxation^39^. DENSE imaging typically employs relatively large voxel sizes (5–16 mm^3^) given the limited SNR of stimulated-echo acquisitions and the substantial SNR needed for strain calculations, which require computation of spatial derivatives. However, focusing on measuring linear displacement improves sensitivity and can therefore accommodate the lower SNR levels that accompany smaller voxel sizes. Larger voxels are inherently more vulnerable to partial-volume effects, such as between tissue and CSF compartments. These partial-volume effects could bias the quantification of tissue motion; however, these potential biases have not yet been fully investigated. Here, we present DENSE tissue motion measurements using various voxel sizes with the goal of improving spatial precision and characterizing possible bias due to partial-volume effects.

For the present study, we focused on the brainstem for multiple reasons: brainstem tissue is expected to yield relatively large amplitude cardiac-locked motion; multiple previously published measurements are available for comparison; and the presence of surrounding fast-moving CSF allows for a characterization of partial-volume effects. Thus, the aims of the study are to introduce and validate a DENSE implementation with retrospective cardiac gating, and to apply it to the quantification of brainstem motion to evaluate its precision and accuracy.

## Methods

### Participants

Eight volunteers (3 females, 24–37 years of age) participated in the study after providing written informed consent following our institutional review board approved protocol and following the guidelines of our hospital. Two volunteers returned for a repeat session.

### MRI acquisition

All data were acquired on a whole-body 7 Tesla Siemens Magnetom Terra (Siemens Healthineers, Erlangen, Germany) at the Athinoula A. Martinos Center for Biomedical Imaging using an inhouse-built 64-channel receive coil brain array for signal reception and a single-channel birdcage volume coil for transmission^40^.

An anatomical 3D multi-echo MPRAGE T_1_-weighted scan^41^ adapted to 7 Tesla^42^ was acquired for definition of brainstem regions with the following parameter values: voxel size = 1 mm isotropic, TR = 3500 ms, TEs = 1.42/2.76/4.1/5.44 ms, TI = 1300 ms, echo spacing = 7.2 ms, non-selective excitation flip angle = 7°, GRAPPA factor = 3.

### DENSE-EPI acquisition

To enable measurement of tissue motion as a function of the cardiac cycle, we implemented DENSE using a single-shot stimulated-echo EPI readout, as described previously^28,32^, with some notable changes: cardiac gating was implemented retrospectively rather than prospectively; single-sided (i.e. single-polarity motion-sensitizing gradients) motion encoding was used for background-phase removal; and a 90° flip angle was used for each measurement instead of variable flip angles throughout the cardiac cycle, resulting in a single, well-defined mixing time (TM).

A schematic of the pulse sequence is provided in Figure 1, which defines parameters TE, TR, TM, G, δ, and Δ. A set of axial slices was centered on the pons for acquisition of the whole brainstem. The pulse sequence is identical to a low-b-value stimulated-echo diffusion acquisition^44^ and consists of a pair of pulsed gradients for motion encoding. Displacement encoding is achieved by modulating the first moment of the motion-encoding gradients akin to phase-contrast techniques. The velocity encoding or *v*_enc_ value is defined conventionally as the maximum velocity measured without aliasing, and is given by

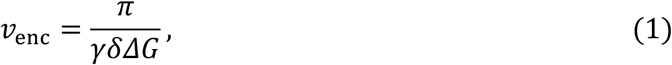

where *γ* is the gyromagnetic ratio and G is the amplitude of the motion-sensitizing gradients. In the context of displacement imaging, an analogous *d*_enc_, or displacement encoding, value can be defined according to

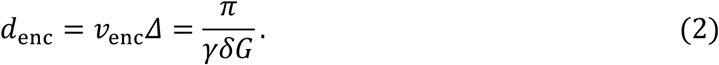

**Figure 1.**
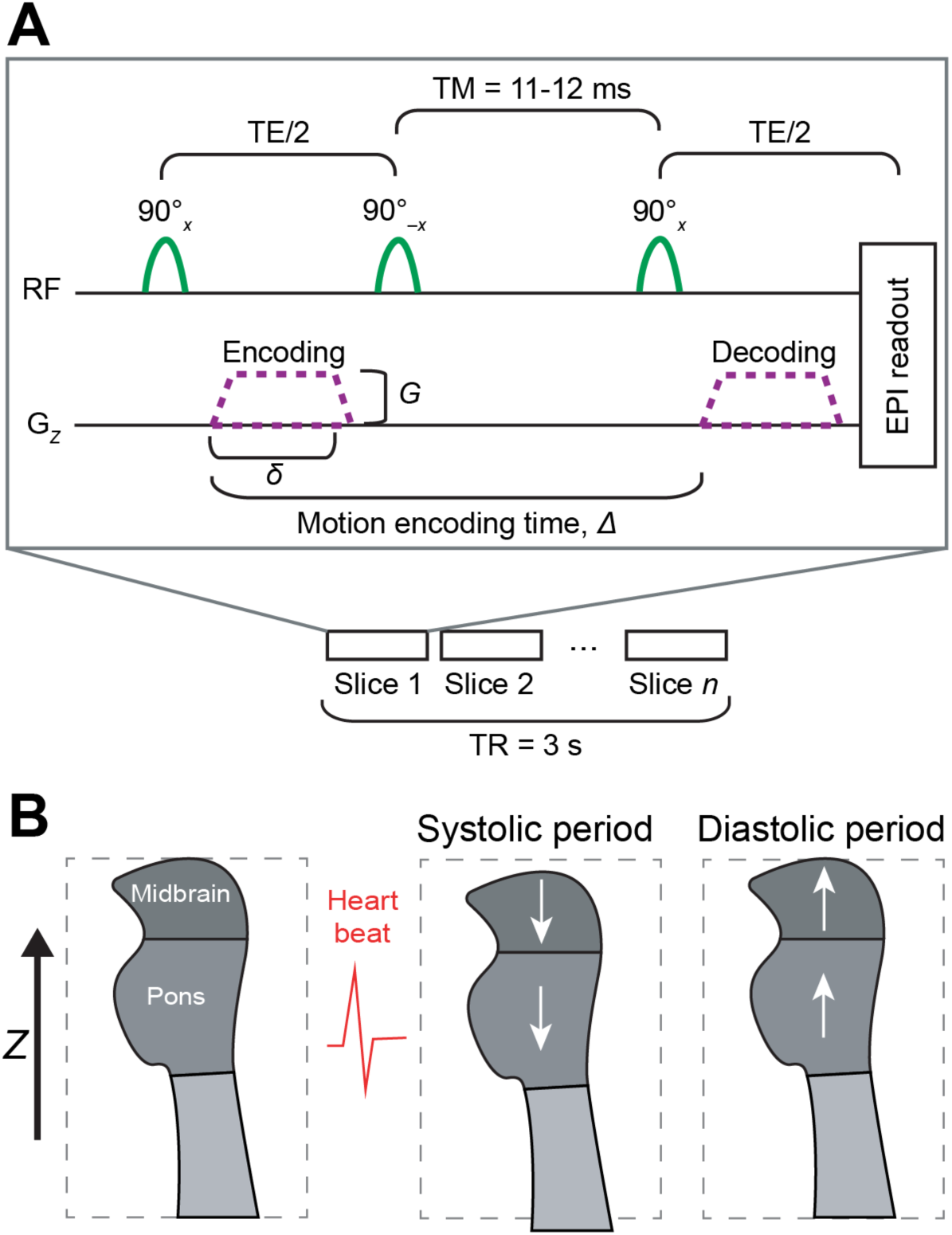
Simplified diagram summarizing DENSE implementation and expected motion pattern. (A) DENSE pulse sequence. Motion encoding was performed in the *Z* direction only. Spoilers and slice select gradients are not shown. Motion encoding is implemented with a pulsed-gradient pair, akin to the pulsed-gradient spin-echo acquisition used in diffusion MRI; however, here the pulse sequence is identical to a STEAM diffusion acquisition. Motion-encoding gradients were toggled on and off between volume acquisitions. Parameter values for each protocol corresponding to the four voxel sizes used are provided in Table 1. TM = mixing time. (B) Expected pattern of midbrain and pons motion over the cardiac cycle based on previous studies. Downward motion relative to the *Z* direction and G*_Z_* gradient is expected during the systolic period (approximately the first third of the cardiac cycle), followed by upward motion during the diastolic period.

**Table 1.** Parameter values for the four acquisition protocols with varying voxel sizes.

| Parameter | $1.2 \times 1.2 \times 2 \text{ mm}^3$ | $2 \times 2 \times 2 \text{ mm}^3$ | $3 \times 3 \times 3 \text{ mm}^3$ | $4 \times 4 \times 4 \text{ mm}^3$ |
| --- | --- | --- | --- | --- |
| TR (ms) | 3000 |  |  |  |
| Temporal resolution (ms) | 6000 |  |  |  |
| TE (ms) | 54 |  |  |  |
| Excitation flip angle | $90^\circ$ | | | |
| Run duration (min) | 2.2 |  |  |  |
| Number of runs | 4 | 2 | 2 | 2 |
| Number of slices | 32 | 32 | 18 | 18 |
| FOV ($\text{mm}^2$) | $192 \times 192$ | $256 \times 256$ | $192 \times 192$ | $256 \times 256$ |
| Nominal echo spacing (ms) | 0.71 | 0.53 | 0.53 | 0.53 |
| GRAPPA factor | 4 | 3 | 2 | 2 |
| TM (ms) | 12 | 12 | 12 | 11 |
| $v_{\text{enc}}$ (mm/s) | 1.8 | 1.8–2.4 | 3.1–4.3 | 3.2 |
| $\delta$ (ms) | 7.3 | 9.9 | 13.2 | 13.2 |
| $\Delta$ (ms) | 37.6 | 38.2 | 38.5 | 37.5 |
| G (mT/m) | 23.7 | 13.0–17.6 | 5.4–7.4 | 7.3 |
| b-value ($\text{s}/\text{mm}^2$) | 75 | 41–75 | 12–23 | 22 |

The *d*_enc_ is independent of TM and is thus particularly useful for acquisitions with multiple TM values. Here, we use a single TM and opt for the native metric of velocity rather than displacement. Note that velocity estimates can be converted to displacement estimates by temporal integration.

Motion-encoded frames (Phase*_Z_*) were interleaved with non-motion-encoded frames (Phase_0_) for subtraction of background-phase offsets. Because background phase may vary over time within each run, this scheme enabled the isolation of phase changes induced by the motion-sensitizing gradients. (It was later determined that this regular sampling of the background phase is not necessary for our stimulated-echo EPI, and can be less frequent to increase temporal efficiency.) Coil-channel-dependent phase offsets were corrected online using ASPIRE^45^ to generate phase-valued images without singularities.

Data were acquired with four voxel sizes to test for the influence of partial-volume and SNR on tissue motion estimates: 1.2×1.2×2 mm^3^, 2×2×2 mm^3^, 3×3×3 mm^3^, and 4×4×4 mm^3^. The protocols used for each voxel size are provided in Table 1. Motion-sensitizing gradients were applied only in the *z* direction in scanner coordinates, which corresponded to the Head-Foot direction. Each run consisted of 40 repetitions, i.e., 20 Phase*_Z_*–Phase_0_ pairs, yielding 20 motion-encoded volumes per run. The TR, TE, and excitation flip angle values were held constant. Inadvertent differences in the *v*_enc_ value between the four protocols were caused by small errors in the original *v*_enc_ calculations discovered after data acquisition.

Cardiac waveforms were recorded concurrently with the DENSE acquisition using an external piezoelectric device on the fingertip.

### Preprocessing

The preprocessing pipeline is summarized in Figure 2. Contributions to phase-valued data in a motion-sensitized acquisition include: (a) background-phase from the B_0_ field and from B_1_^+^ and B_1_^−^ fields caused by susceptibility effects, inhomogeneities in the main magnetic field, and the transmit and receive radiofrequency coils; (b) eddy current-induced phase from the motion-sensitizing gradients; and (c) the desired phase shift from moving spins due to deliberate motion encoding provided by the motion-sensitizing gradients.

**Figure 2.**
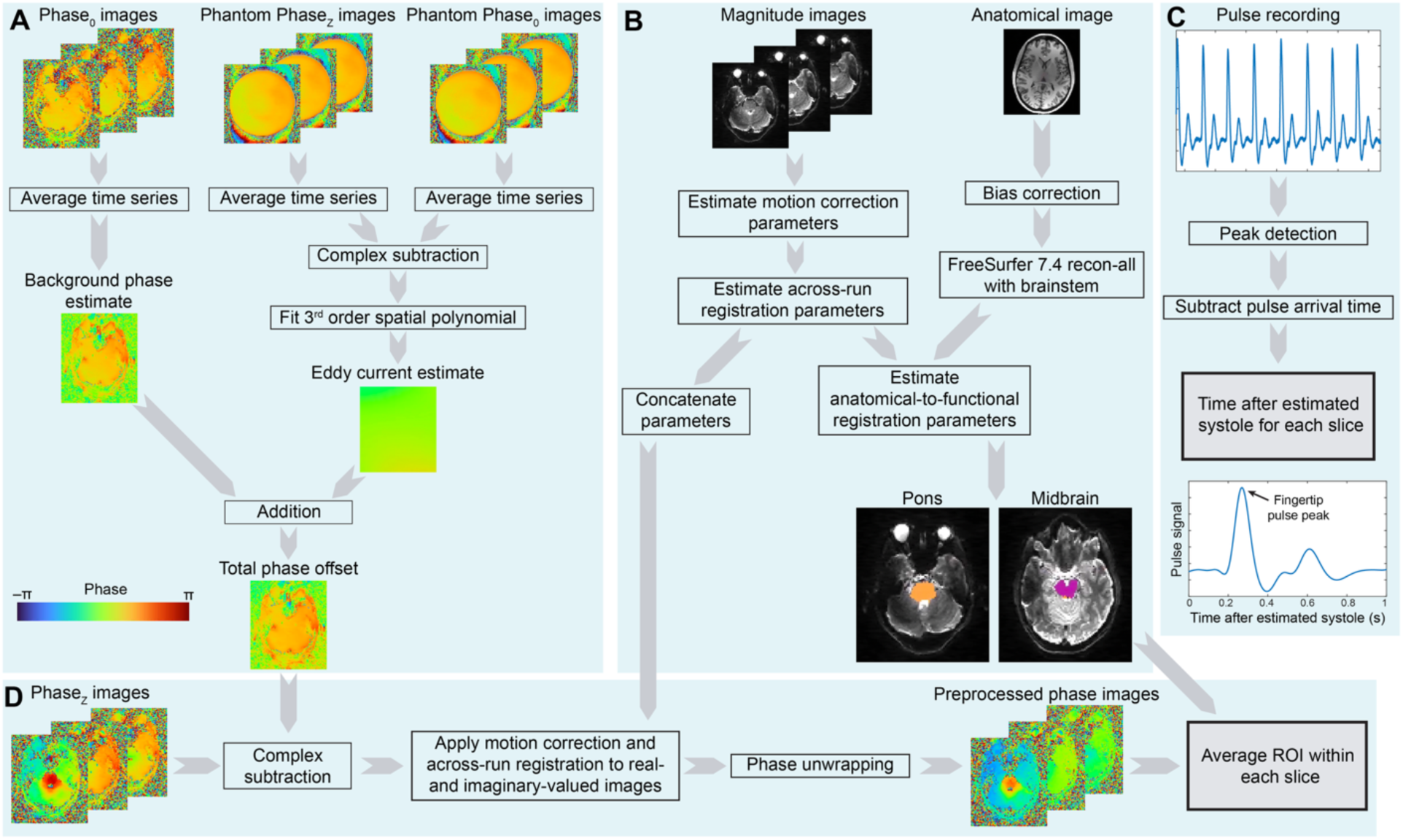
Illustration of processing pipeline for cardiac-related velocity in the brainstem. Pipeline is repeated for each voxel size protocol separately. Time after systole and average velocity within each brainstem ROI become the horizontal and vertical axes, respectively, in Figure 3. (A) Estimation of phase offset including background-phase and eddy-current contributions. Static phantom images are acquired in a separate session with matched slice prescription and acquisition parameters. (B) Procedure for motion correction, across-run registration, and ROI selection. (C) Summary of the processing of pulse device recordings to estimate time after systole for each slice and time point, including an average time course of the pulse device signal as a function of the absolute time after estimated systole. (D) Summary of processing for motion-encoded *in vivo* phase images, applying information from panels A and B. See text for further details. Phase*_Z_* = motion-encoded phase, Phase_0_ = non-motion-encoded phase, ROI = region of interest.

To correct for background-phase contributions (Figure 2A), volumes without motion encoding (Phase_0_) were first averaged together within each run to generate a background-phase estimate. To correct for eddy current-induced phase present only in the motion-encoded frames (Phase*_Z_*), data were acquired for each of the four protocols in a separate session using a spherical agar phantom with slice prescriptions that were identical to those of the *in vivo* acquisitions; this enabled estimation of the slice-prescription-dependent eddy current-induced phase. To extract eddy-current-induced phase, we first corrected for background-phase offsets identically to the *in vivo* pipeline, then masked the image based on magnitude-valued image intensity, and finally fit a third-order spatial polynomial within each slice. This smoothly varying eddy-current-phase estimate was combined with the subject-specific background-phase estimate for subtraction of confounding phase contributions.

Phase wraps were present in some slices in the high spatial resolution data (1.2×1.2×2 mm^3^ and 2×2×2 mm^3^) due to the higher velocity sensitivity (i.e., the lower *v*_enc_ value) used. Because these data were relatively noisy, applying phase unwrapping to every slice sometimes resulted in inappropriate unwrapping. Therefore, slices that required phase unwrapping were identified and labelled manually to apply the unwrapping algorithm only to slices and timepoints that required unwrapping (Figure S1). To avoid unnecessarily unwrapping phase in the noise regions outside of the brain, images were masked with a brain mask defined on the anatomical image with the FSL tool *bet*^46^. Automated phase unwrapping was then applied in the identified slices using a 2D algorithm^47^.

Motion estimation and across-run registration (Figure 2B) were performed using the magnitude-valued data with *mri_robust_template* from FreeSurfer. For resampling, the complex-valued images were first separated into real and imaginary components then resampled to a template that was common across runs, or an across-run template. One across-run template was defined for each subject and voxel size protocol. Since our DENSE-EPI acquisition is a 2D multiple-slice method, each slice within each volume was acquired at a specific time within the cardiac cycle. Given that our data exhibited through-slice motion, to preserve the acquisition time for each voxel and thus the time within the cardiac cycle, we resampled the images with nearest neighbor interpolation. Similarly, to track the location of each slice and thus the precise cardiac timing of each voxel, we created a synthetic slice-index volume that was also resampled to template space using nearest-neighbor interpolation to retain the integer-valued indexing. All analyses were completed in the across-run template space corresponding to the specific protocol.

### Analysis of cardiac-locked tissue motion

First, the time series recorded from the pulse device was temporally low-pass filtered. Then, the *findpeaks* function in MATLAB (Mathworks, Natick, MA) was used to identify pulse peaks. An average pulse arrival time of 269 ms based on literature values^41^ was subtracted from the time of each pulse peak to account for the pulse arrival time from the heart to the fingertip, so that each identified peak corresponded to an estimated time of cardiac depolarization, or the R wave on an electrocardiogram, which we will refer to as the “time of estimated systole”. Finally, we generated the cardiac response as a function of the absolute time after systole (Figure 2C). This approach differs from the common approach of characterizing the cardiac response as function of the relative time within the cardiac cycle, i.e., cardiac phase^48^. To generate the cardiac response, the absolute time after estimated systole (in units of ms) was calculated for each slice in the DENSE-EPI acquisition.

We calculated the cardiac-locked tissue velocity by using the median velocity within 25-ms time bins. Heart rate varied between and within individual subjects, so an arbitrary cutoff of 1 s was used as the end of the cardiac cycle. The mean velocity across bins was subtracted from each subject- and voxel size-specific velocity time course. To compare our quantifications with previously reported peak displacement values, we calculated the cardiac-locked tissue displacement by cumulative integration over the cardiac-locked velocity time course and subsequently selected the maximum displacement value for each subject and voxel size.

We used a Wilcoxon signed-rank test with Bonferroni correction to test whether velocity measurements within each cardiac bin were significantly different than zero. To assess differences in cardiac-locked velocity measurements across voxel sizes and subjects, we used a two-way ANOVA, with voxel size treated as an ordinal variable. To evaluate uncertainty in displacement measurements, subject-level standard error (for between-subject analyses) or 95% confidence intervals (for within-subject analyses) were estimated using non-parametric bootstrapping with 1000 iterations.

### Region of interest definition

To process the anatomical reference data, the T_1_-weighted images were bias-field corrected^49^ before automatic brain segmentation and cerebral cortical surface reconstruction were performed using FreeSurfer *recon-all*. The pons and midbrain were identified on anatomical images with the brainstem segmentation tool in FreeSurfer^50,51^. Rigid (6 DOF) registration parameters between DENSE-EPI across-run templates and the anatomical reference were estimated with the FreeSurfer tool *bbregister*^52–54^, then applied to the pons and midbrain masks to register them to each of the DENSE-EPI across-run templates. These anatomical masks provided regions of interest (ROIs) for subsequent analysis of the DENSE-EPI data. Spatial averaging was achieved by selecting the median phase value within slices of the ROIs.

### Flow phantom validation

To characterize the quantitative accuracy of our velocity measurements, we performed DENSE-EPI on a slow-flow phantom that produced water flow with a known velocity using a calibrated syringe-based pump (Pump 11 Elite, Harvard Apparatus, Holliston, MA, USA), similar to previous work^55–57^. Flow velocity accuracy was confirmed separately by recording the rate of filling of a graduated cylinder at the exit port of the tube (data not shown). Tap water was directed through flexible tubing with an inner diameter of 6.4 mm into the MRI scanner bore. The tubing was wrapped around an agar gel-filled bottle such that four axial cross sections of the tube were visible in the acquired volume. We varied the water flow velocity setting of the pump (0.1, 0.2, 0.4, 0.8, and 1.5 mm/s) and *v*_enc_ value of the acquisition (1, 2, and 3.5 mm/s) using only the 2×2×2 mm^3^ protocol to assess bias. We then acquired all four voxel size protocols with the flow velocity set to 0.5 mm/s. All acquisitions were repeated with the pump turned off to generate a velocity of 0 mm/s (“flow-off images”). For analysis, flow-on images were divided by the flow-off images using the complex-valued image reconstructions, equivalent to phase subtraction, to correct for confounding phase contributions. ROIs centered on the cross-sections of the tube were created with a magnitude-valued signal threshold. Phase values were averaged across all four tube cross-sections after accounting for direction of flow.

### Simulation of partial-volume effects on velocity estimation

The complex-valued signal within a voxel containing CSF and tissue contributions can be expressed as a linear combination of complex-valued components:

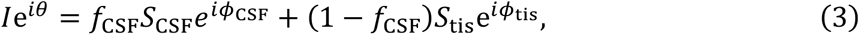

where *I* and *θ* are the signal intensity and phase, respectively, of the voxel, *f*_CSF_ is the partial-volume fraction of CSF, *S*_CSF_ and *S*_tis_ are the signal intensities of CSF and tissue, respectively, and *ϕ*_CSF_ and *ϕ*_tis_ are the signal phases of CSF and tissue compartments in the voxel, respectively. Tissue velocity was fixed at 1 mm/s, tissue signal intensity was fixed at 20, and a *v*_enc_ value of 3 mm/s was used. Simulations were implemented in MATLAB.

## Results

### Brainstem motion across the cardiac cycle

We successfully acquired tissue velocity measurements in all eight participants, and two repeat sessions. Figure 3A depicts cardiac-locked velocity patterns for each voxel size in the pons and midbrain. We observed a consistent double-peaked velocity waveform, with two distinct peaks of head-to-foot directed velocity detected for all voxel sizes at approximately 125–150 ms and 275–300 ms after estimated systole. Pooling across voxel sizes and subjects, binned measurements were significantly different than zero for most time bins after Bonferroni correction (Fig. 3A). There were no significant differences across voxel sizes for either velocity peak when controlling for differences across subjects (first peak for pons: *F*(3,21) = 2.3, *p* = 0.10; second peak for pons: *F*(3,21) = 0.6, *p* = 0.62; first peak for midbrain: *F*(3,21) = 1.1, *p* = 0.37; second peak for midbrain: *F*(3,21) = 1.9, *p* = 0.15).

**Figure 3.**
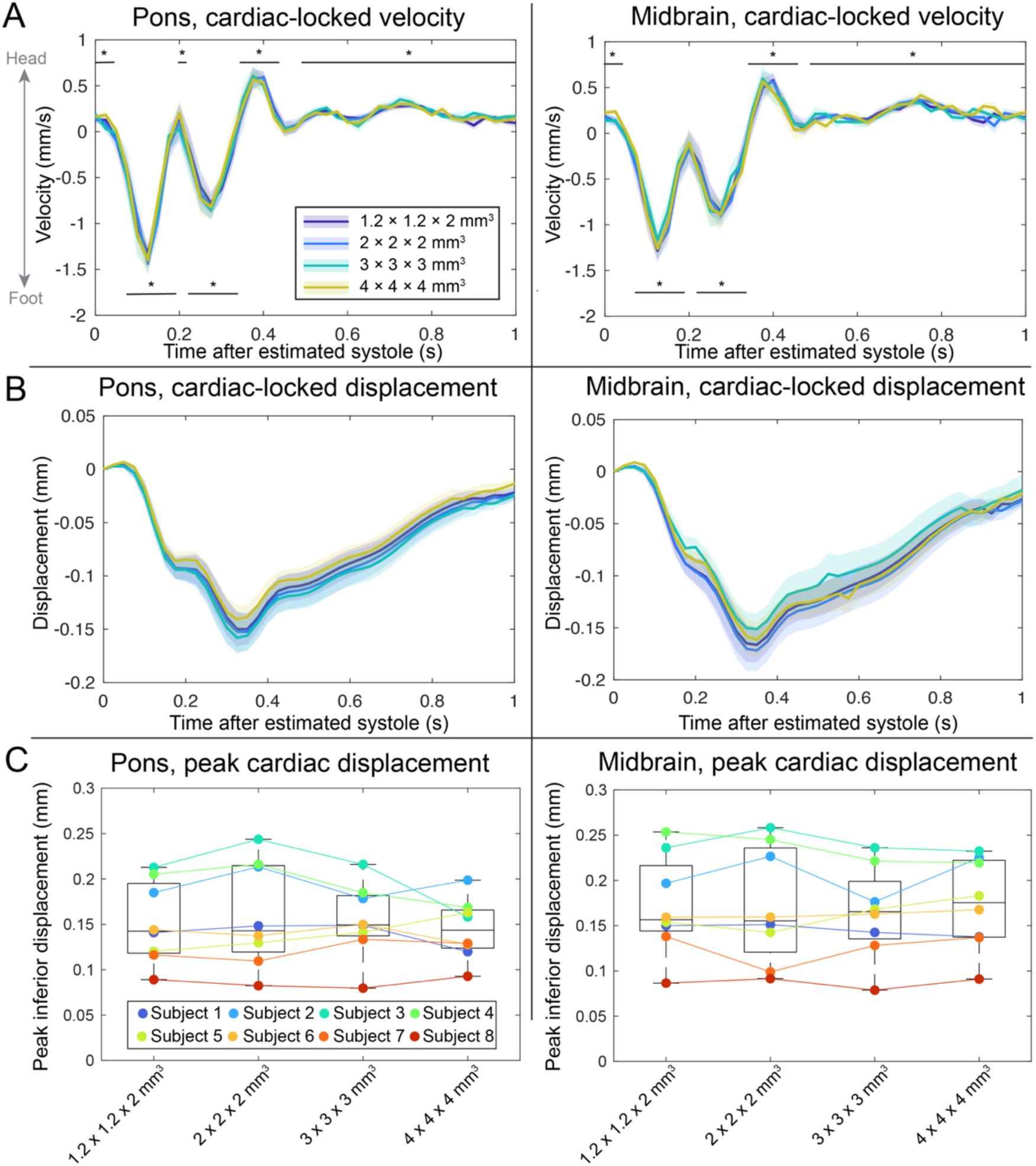
A double-peaked waveform is detected in pons and midbrain, across voxel sizes. (A) Native velocity measurement within each ROI, averaged within 25-ms bins across subjects. Asterisk (*) indicates *p*<0.05 using Wilcoxon signed-rank test, Bonferroni corrected. Error bars indicate standard error across subjects. (B) Cardiac-locked velocity responses from panel A were cumulatively integrated over time to calculate cardiac-locked tissue displacement. Error bars indicate standard error for each cardiac bin estimated using nonparametric bootstrapping with subject-level resampling. (C) Peak cardiac displacements for each subject and voxel size protocol derived from time-integrated displacement calculations. Color indicates individual subjects. Positive velocity corresponds to upwards foot-to-head directed motion.

Cumulative integration of the velocity responses revealed average peak displacements of 0.150±0.016 mm for the pons and 0.163±0.017 mm for the midbrain (Figure 3B). Peak displacements did not significantly differ across voxel sizes when controlling for differences across subjects (pons: *F*(3,21) = 1.3, *p* = 0.29; midbrain: *F*(3,21) = 0.9, *p* = 0.47; Figure 3C).

To assess whether the definition of the cardiac cycle timing in terms of absolute time or relative time impacts the resulting velocity responses, we temporally resampled our velocity time courses as a function of cardiac phase and compared them to our proposed absolute time after systole metric. Re-binning as a function of cardiac phase obscured the double-peak velocity pattern (Figure 4A). This was primarily due to averaging across subject-specific time courses with different underlying heart rates (Figure 4B). However, the cancellation of the double-peak pattern also occurred in some individual subjects with high heart rate variability (Figure 4C). In contrast, the double-peak velocity pattern was detected in all individual subjects when using absolute time after estimated systole.

**Figure 4.**
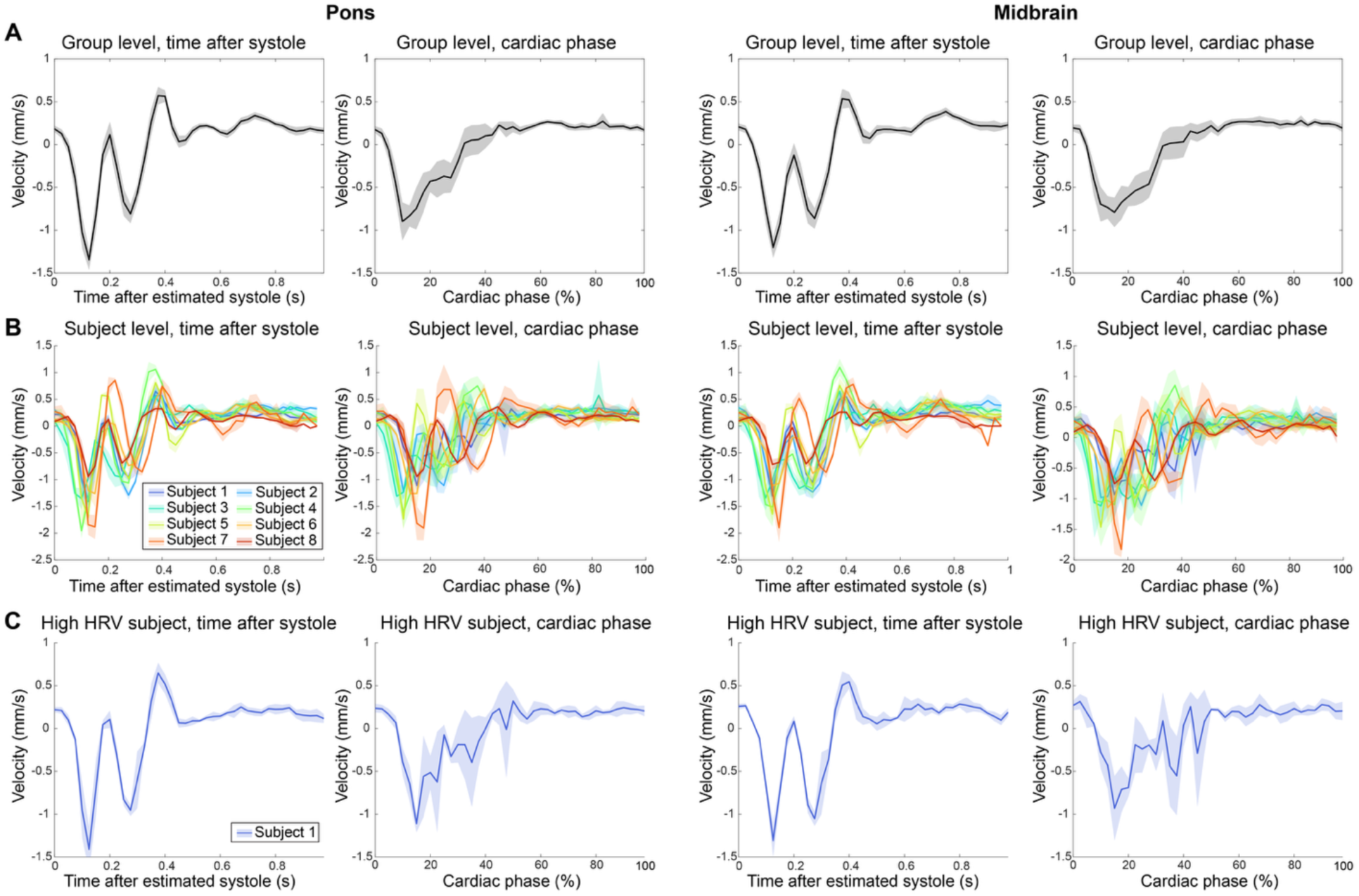
Velocity dynamics detected with absolute time after systole can be obscured when analyzed using cardiac phase. (A) Cardiac-locked velocity responses averaged across subjects and voxel size, binned according to time after systole (as in Figure 3A) or cardiac phase (percentage of the cardiac cycle). The origin of the time axis (i.e. time *t*=0) is the time of estimated systole for both. Error bars indicate standard error across subjects. (B) Single-subject cardiac-locked velocity time courses after averaging across voxel sizes. Error bars indicate standard deviation across voxel size protocols. (C) Example subject from panel B (Subject 1) with especially high Heart Rate Variability (HRV), plotted independently to visualize possible discrepancy between the two methods to compute cardiac response over time for an individual subject. Error bars indicate standard deviation across voxel size protocols.

### Precision and accuracy assessed via scan-rescan repeatability and slow-ffow phantom

To assess the precision of our measurement, we acquired repeat acquisitions in two subjects, separated by either 4 or 8 months. The cardiac-locked velocity responses for each session are depicted in Figure 5 and demonstrate robust repeatability of the double-peak velocity waveform in both subjects. Peak displacement measurements between sessions differed by 0.5–6.2% (95% confidence intervals within ±15%).

**Figure 5.**
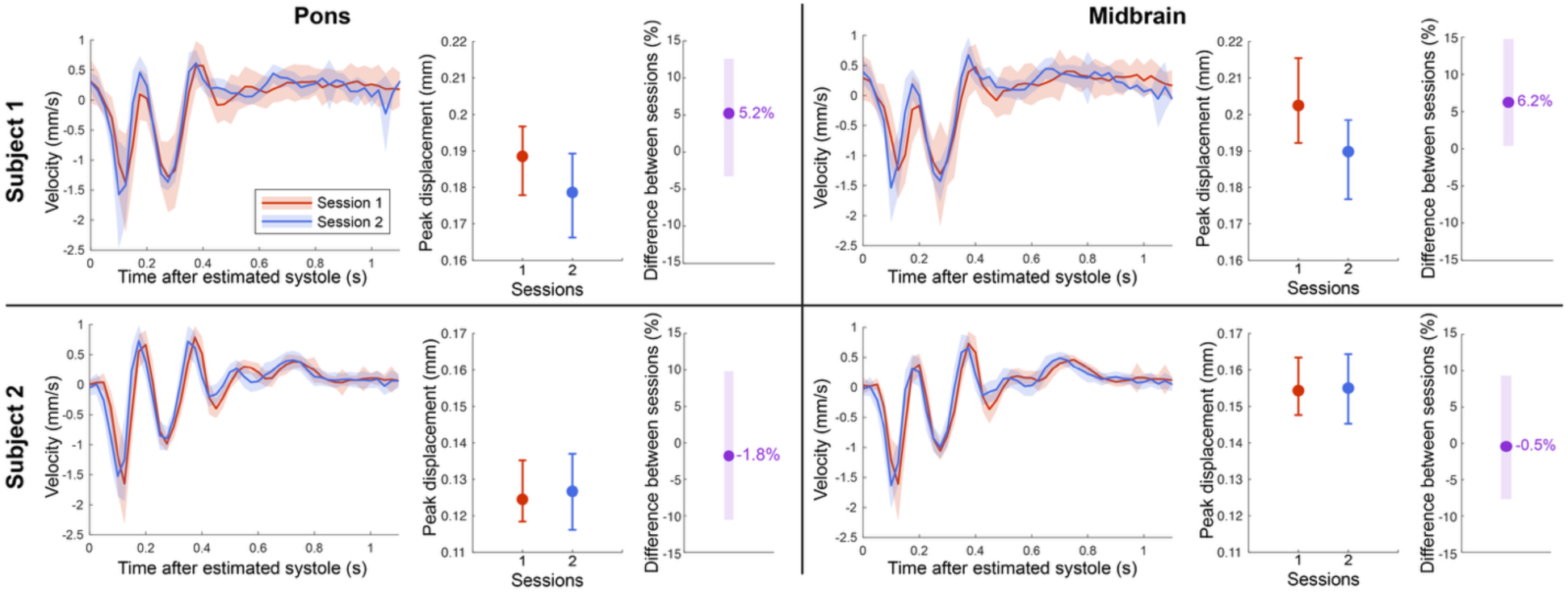
Repeated measurements in two subjects demonstrating scan-rescan consistency. Single-subject cardiac-locked velocity time courses within each ROI, analogous to Figure 3A, taken from two experimental sessions across different days. Measurements from different spatial voxel sizes are averaged together to compare across repeat sessions. Error bars for velocity responses indicate standard deviation within each bin. Peak displacement estimates from each session (red/blue), and the percent difference between sessions (purple), are plotted to the right of each velocity response plot, where error bars indicate the 95% confidence interval based on nonparametric bootstrapping. Positive velocity corresponds to upwards foot-to-head directed motion.

To assess the accuracy of our measurement, we repeated our DENSE-EPI measurements in the slow-flow phantom. We observed a bias where larger flow velocities resulted in greater underestimation of velocity, but no differences across *v*_enc_ (*F*(1,9) = 25.3, *p* < 0.001 for velocity; *F*(1,9) = 0.6, *p* = 0.47 for v_enc_; Figure 6B). Flow phantom measurements across voxel sizes showed a bias where larger voxels yielded greater velocity estimates, but this effect is explained by spatial averaging across a laminar flow profile and is not relevant for the *in vivo* tissue displacement measurements (Figure S2).

**Figure 6.**
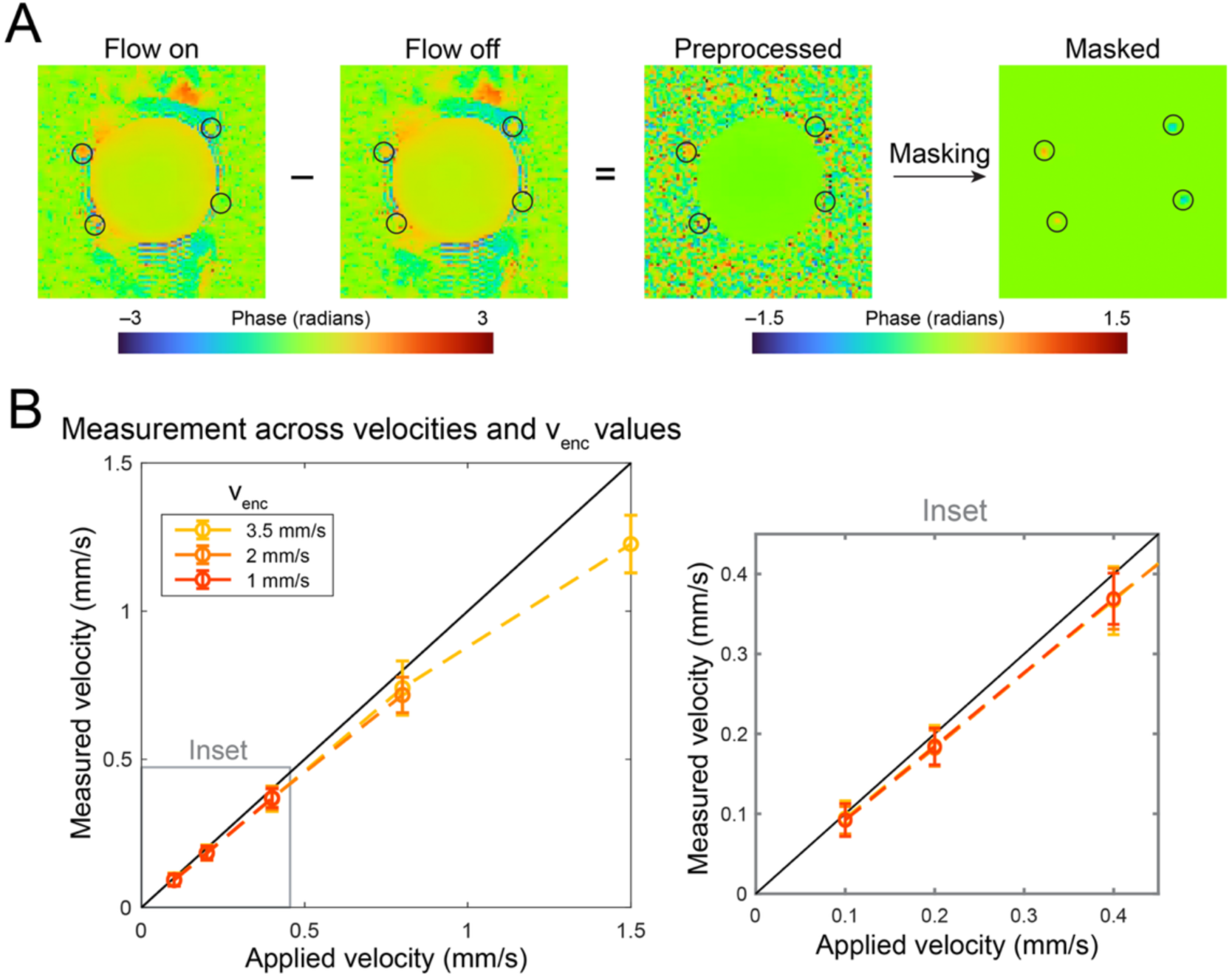
Validation based on slow-flow phantom. (A) Preprocessing pipeline for flow phantom images. Complex-valued images acquired when the flow pump is turned on (“flow on” images) are divided by the complex-valued images acquired without flow (“flow off” images), and the phase of the ratio is taken as the phase-difference image. The stationary agar phantom is centered within the image, and the four tubes are indicated with black circles. The resulting images are masked to only include phase values in the tubes. (B) Measurements for varied applied velocities and *v*_enc_ values. The black line indicates a perfect correspondence between applied and measured flow velocities. Inflow and outflow tubes are combined after reversing the sign of outflow phase values. Error bars indicate standard deviation across slices and time points.

### Partial-volume effects between brainstem and CSF

To investigate the contribution of partial-volume effects to our velocity measurements, we first created ROIs of CSF-contaminated voxels, defined as the edge voxels of our original pons and midbrain ROIs. Cardiac-locked velocity patterns measured with the edge voxels were indistinguishable from velocity patterns derived from the full anatomical ROIs (Figure S3).

To investigate why CSF partial-volume effects did not bias the tissue measurements, we simulated the influence of partial-volume effects between the brainstem and surrounding CSF using CSF velocities reported in the literature^55^. We found that the lack of an effect of voxel size on brainstem velocity measurements could be explained by the large CSF velocities relative to our *v*_enc_ value combined with a large within-voxel variability in CSF velocity, or a wide velocity distribution of neighboring CSF (Figure 7).

**Figure 7.**
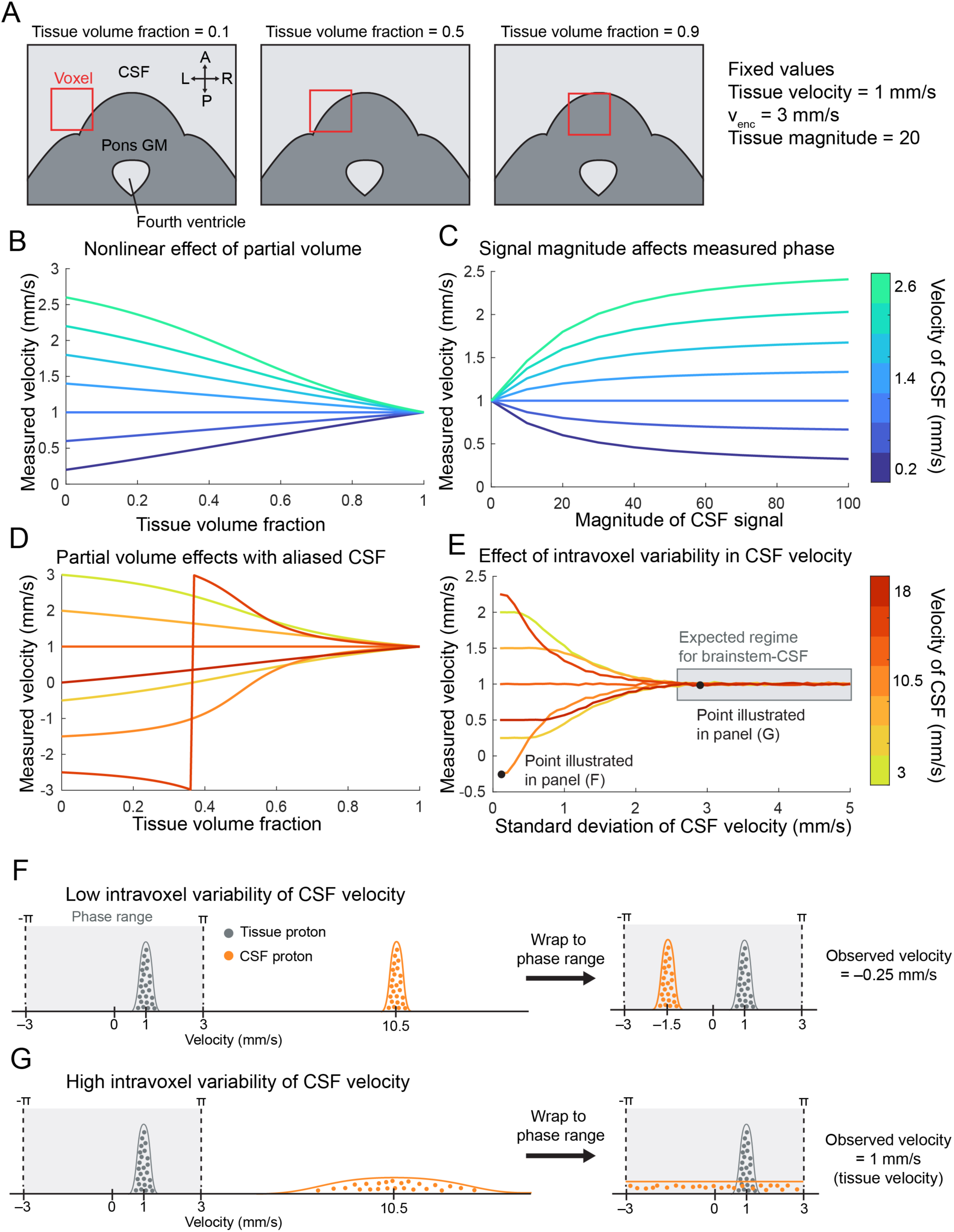
Simulation of expected partial-volume effects on phase-based velocity measurements. (A) Illustration of simulated voxel sampling both gray matter (GM) tissue and CSF with varying partial volumes of each based on Equation 3. The simulations assume a fixed tissue velocity, acquisition *v*_enc_ value, and tissue magnitude-valued image intensity. (B) Measured velocity from image phase as a function of signal fraction of tissue. CSF magnitude-valued image intensity = 20 (i.e., equal to tissue), within-voxel variability in CSF velocity = 0. (C) Measured velocity as a function of magnitude-valued image intensity of CSF. Tissue volume fraction = 0.5, within-voxel variability in CSF velocity = 0 mm/s. (D) Illustration of the effect of aliasing of CSF velocities on measured velocity as a function of tissue volume fraction. CSF magnitude-valued image intensity = 20 (i.e., equal to tissue), within-voxel variability in CSF velocity = 0. (E) Effect of within-voxel variability in CSF velocity (i.e., the spread of the CSF velocity distribution within the voxel) on measured velocity. The highlighted region represents the expected relationship between tissue and CSF velocities near the brainstem. Tissue volume fraction = 0.5, CSF magnitude-valued image intensity = 20 (i.e., equal to tissue). (F–G) Schematic illustrating distribution of proton phases within a voxel in the scenario with low (F) and high (G) within-voxel variability of velocity plotted in panel E, using a mean CSF velocity of 10.5 mm/s as an example. Histograms on the left side represent the true velocities present within the simulated voxel, whereas histograms on the right side represent the observable velocities within the phase range defined by the defined acquisition *v*_enc_ value.

### The double-peak pattern reffects two separable sources of motion

The two head-to-foot-directed velocity peaks that we observed in the pons and midbrain had distinct spatial patterns. The first peak was local, restricted primarily to the brainstem itself, whereas the second peak was global, encompassing all structures seen within an axial slice (Figure 8A). To provide a quantitative assessment of this observation, we plotted one-dimensional profiles of the average cardiac-locked velocity response along the Anterior–Posterior and Left–Right directions centered on either the pons or midbrain at the time point corresponding to either the first velocity peak or second velocity peak. These one-dimensional profiles confirm that there was a nonuniform spatial profile of the velocity response at the first peak that is maximal within the brainstem, and a more uniform spatial profile at the second peak (Figure 8B).

**Figure 8.**
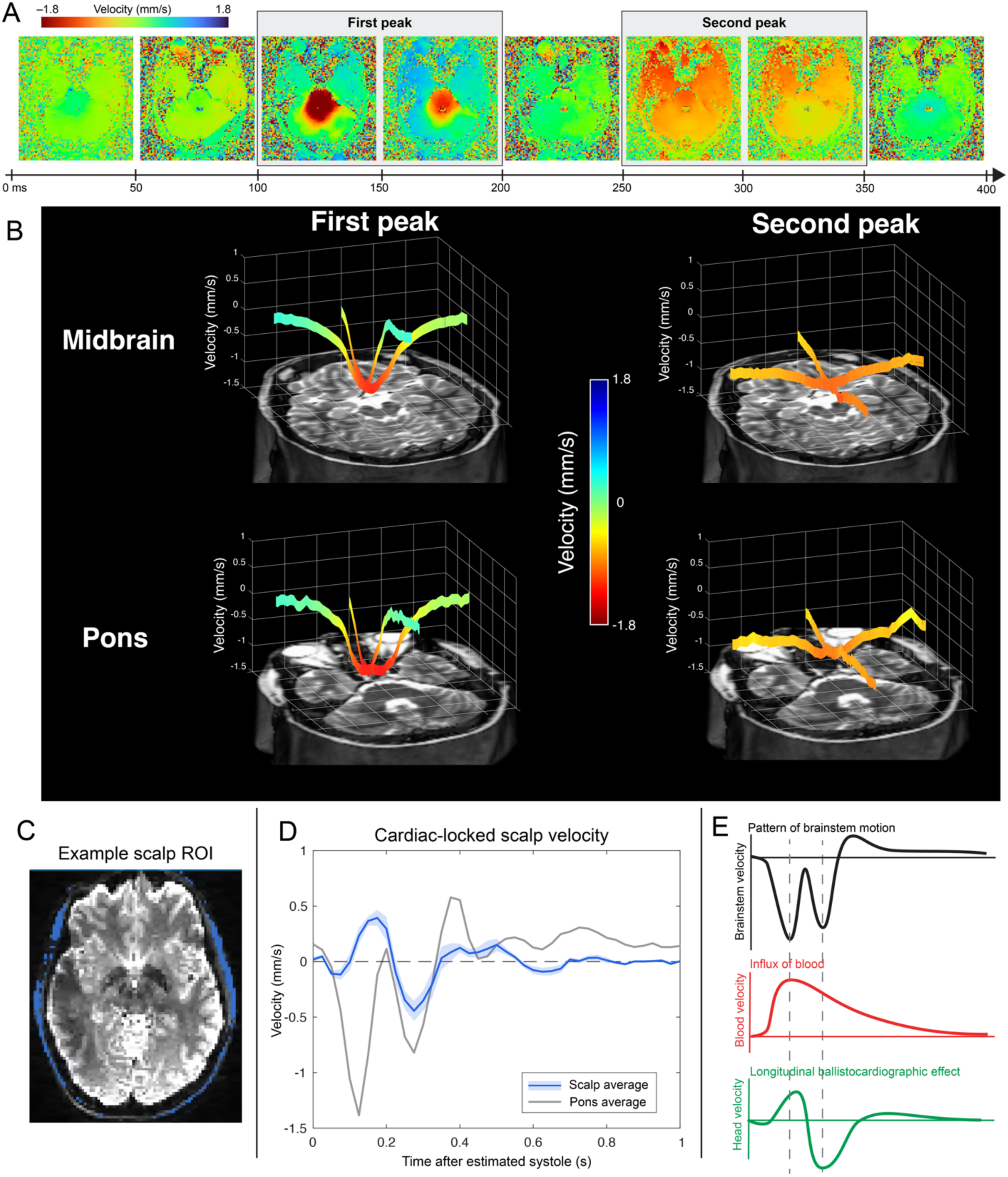
Two distinct spatial patterns within the double-peak pattern in cardiac-locked velocity response. (A) Whole-slice velocity averages within 50-ms bins in a representative subject using the 2-mm isotropic voxel size over the first 400 ms of the cardiac cycle. The origin of the time axis (i.e. time *t*=0 ms) refers to estimated systole. (B) Spatial profiles of motion for each velocity peak. Profiles are averaged within a column of tissue in either the Anterior–Posterior or Left–Right directions, as indicated by the underlying anatomical volumetric renderings. Data are averaged across subjects and slices and include only the 2-mm isotropic voxel size. (C) Example scalp ROI for each imaging voxel size used to create the traces plotted in panel D in a single representative subject. (D) Scalp ROI cardiac-locked velocity time courses averaged over voxel sizes. The average pons velocity time course is plotted in gray for comparison. (E) Schematic representing hypothesized time courses of the approximate pattern and timing of Peak 1 and Peak 2 of brainstem motion compared to the influx of blood to the cranium and longitudinal rigid motion of the head^35^, respectively.

To investigate whether the uniform spatial pattern of the second peak is the result of global, perhaps rigid head motion, we sought an ROI that is expected to move rigidly along with the skull and not affected by brain tissue deformation. For this, we chose ROIs within the scalp to estimate contribution from well-known cardiac-locked rigid head motion identified in previous studies and in cardioballistography^51^. The scalp ROIs were created for each voxel size using a semi-manual approach based on concentric ellipses (Figure 8C). We observed a bipolar pattern of cardiac-locked velocity in the scalp with a peak of head-to-foot-directed motion that coincided in time with the second peak observed in the brainstem motion time courses (Figure 8D).

## Discussion

We present an implementation of the DENSE method that revealed a thus far unreported double-peak pattern of cardiac-locked brainstem velocity. We were able to achieve the smallest voxel size to date for DENSE (2.88 mm^3^) due to the SNR advantages at 7T and because we chose to not calculate strain metrics. Since we used a single TM value, we report tissue motion in units of velocity, rather than displacement. We used four different protocols with increasing voxel size and hypothesized that velocity estimates would vary with voxel size; however, we found negligible bias in velocity estimates with voxel size.

The precision of our measurement is relatively high, as evidenced by the high repeatability across sessions. Peak cardiac-locked displacement measurements across sessions varied by 0.5–6.2%, compared to approximately 20% reported in a previous study using the DENSE method, though we measured tissue motion in a different brain region^32^. We maintained temporal precision in part by carefully tracking the acquisition timing of each slice within the multiple-slice volume during preprocessing for accurate cardiac timing. The accuracy of our method can be assessed by comparing our measurements to those of previous studies and by the comparison of our measured velocities to ground-truth velocities provided by our flow phantom. Our average peak cardiac-locked displacement estimates were 0.145 and 0.164 mm for the pons and midbrain, respectively, which agree well with previous quantifications of 0.15 to 0.18 mm in these regions^31,32^. There was substantial inter-individual variability, with peak displacements ranging between 0.09–0.22 mm across subjects. In our flow phantom study, we identified a bias in velocity estimation, where faster flow velocities resulted in a greater underestimation of velocity, even after confirming the accuracy of the flow phantom itself. This bias is similar to one reported previously using phase-contrast MRI^55^. However, another study comparing displacement measurements using the DENSE method to a displacement phantom did not report an underestimation of motion^56,57^. If the bias observed in the phantom is also present in our *in vivo* data, then the true average peak cardiac-locked displacement may be somewhat higher. Further work will be required to determine whether this effect is specific to our flow phantom and whether it is relevant for *in vivo* measurements.

We report our cardiac-locked time courses as a function of absolute time after estimated systole, in contrast to the convention of defining them as a function of cardiac phase. This decision was fundamental to our observation of the double-peak pattern of cardiac-locked brainstem velocity, since the re-binning of our data according to cardiac phase obscured this pattern. Physiologically, the source of cardiac-locked brainstem motion is the mechanical energy initiated at cardiac contraction, and its timing is primarily dictated by the propagation time of blood from the heart to the cranium. In contrast, analyzing data as a function of cardiac phase assumes that the phenomenon of interest occurs at a fixed percentage of the cardiac cycle, regardless of the time delay from the heartbeat. As a result, we found that when binning by cardiac phase, the double-peak velocity pattern became obscured when heart rate varied, both between and within individuals (Figure 4B–C). Ultimately, the decision to define the cardiac response using absolute time or relative time will depend on the hypothesized mechanism by which the cardiac dynamics affect the signal of interest.

We identified a double-peak pattern of cardiac-locked brainstem velocity that has not yet been described, though it can be observed in a previous study^58^. During the first peak, the motion appears spatially restricted to the brainstem, consistent with a nonrigid motion pattern. During the second peak, we observed a more global pattern of motion that coincides with extracranial and presumably rigid cardiac-locked motion in the scalp. We theorize that the first peak in velocity is consistent with the Monro-Kellie doctrine, where arrival of blood volume into the cranium requires the expulsion of CSF and the nonrigid deformation of the tissue through the Foramen Magnum. The timing of the first peak agrees with this hypothesis, as the blood arrives to the head approximately 126 ms after systole^59^, and the first peak we identified occurs 125–150 ms after systole (Figure 8E). Note, however, that our reported cardiac timing is based on an estimated time of systole based on our fingertip pulse recording, and inaccuracy on the order of 50 ms is not unexpected. The second peak may be consistent with cardiac-locked longitudinal motion (Figure 8E). Cardiac-locked rigid head motion has been observed with ballistocardiograms^60^, MRI-compatible camera systems^61^, conventional video^52–54^, EEG electrodes^54^, wearable devices^50,62^, and our own data based on fast single-slice imaging and retrospective motion correction^63^. The pattern of motion we observed in the scalp has both a positive (upwards motion) and negative (downwards motion) lobe, similar to the patterns observed using the independent measures mentioned above, and the second peak in our brainstem cardiac-locked time course coincides with the negative lobe. The precise spatial pattern of this rigid motion is unclear, but its amplitude is likely different between our scalp ROI near the center of the cranium and the brainstem at the base of the skull, so we did not attempt to correct our brainstem data for this presumably overlying rigid motion. Future work aiming to quantify only the nonrigid tissue motion components may benefit from further investigating, and accounting for, this effect.

Unexpectedly, we did not observe an effect of voxel size on our velocity estimates, even in carefully-selected edge voxels that maximized the partial-volume effects between the brainstem and surrounding CSF. Partial-volume effects in phase-valued data depend on the signal magnitude and the presence of aliasing. For instance, our simulations illustrate the nonlinear relationship between CSF signal magnitude and measured velocity in the setting of a fixed partial-volume fraction (Figure 7C). Our simulation results also reproduced the negligible voxel size bias through a combination of (a) phase aliasing, and (b) a broad distribution of velocity (or high intra-voxel variability), within the CSF compartment. Our acquired *v*_enc_ values of 1.8–4.3 mm/s were much lower than the peak velocity of surrounding CSF—approximately 20–35 mm/s in the pontine cistern^64^. Thus, the phase of these CSF spins will likely be aliased multiple times. We found that the spread of velocities within the CSF compartment must be greater than or equal to the *v*_enc_ value to produce negligible bias. While we do not know the precise velocity variability of CSF at the tissue boundaries, CSF velocity may be increasing rapidly with distance from the tissue boundary^54^, perhaps producing a high within-voxel variability on the order of 1.8–4.3 mm/s. Importantly, these assumptions will likely be invalid in other settings, such as partial-volume effects between cortical tissue and subarachnoid CSF, where tissue and CSF velocities are more similar in amplitude; there, a smaller voxel size will likely be more useful to reduce bias from partial-volume effects.

Recently, methods to estimate slow flow and tissue motion with a spin-echo—rather than stimulated-echo—motion-encoded acquisition have been successful^50,62^. A spin-echo approach provides higher SNR, but a stimulated-echo approach is particularly useful in applications where long motion encoding times are beneficial, because the TM allows for longer motion encoding times while maintaining a relatively short TE and thus high tissue signal^29^. In the current study, we used a single, short mixing time for cardiac sampling, but we plan to extend this method to forms of physiological motion that occurs on longer timescales, such as respiratory motion and motion related to neuronal activity via neurovascular coupling.

Our study has several limitations. First, we used a single motion-encoding direction that was approximately parallel to the anatomical axis of the brainstem, so we could not capture motion components directed away from this axis. Therefore, our velocity and displacement measurements are likely underestimated. Second, we used retrospective cardiac gating, which results in uneven sampling of the cardiac cycle. We chose this approach to eventually extend our method to applications beyond characterizing motion across the cardiac cycle. In contrast, prospective cardiac gating provides uniform, consistent sampling of the cardiac cycle, but impedes the analysis of non-cardiac-related effects. Finally, our method for the correction of eddy-current-induced phase requires an independent phantom session, which can be cumbersome. Previous studies have frequently used two-sided motion encoding^63^ or fit a spatial polynomial to regions of “stationary” signal to capture eddy-current-induced phase^64^. Here, we opt for single-sided motion encoding due to our retrospective gating approach, and we cannot assume that any tissue region is stationary given our high velocity sensitivity.

In conclusion, we quantified cardiac-locked velocity in the pons and midbrain using an implementation of the DENSE method with retrospective cardiac gating and a single mixing time to extend the method to applications beyond cardiac-locked tissue motion. We identified a double-peak pattern of cardiac-locked brainstem velocity that was consistent across subjects and sessions when analyzing our data using absolute time after systole instead of cardiac phase. We also demonstrated the smallest voxel size for DENSE tissue motion quantifications to date, enabling higher spatial specificity for small brain regions and for differentiating tissue and CSF compartments. Next, we plan to extend this method to other tissue regions, primarily cerebral cortex, and other drivers of tissue motion, including respiration and neurovascular coupling^29,65^. Measuring tissue motion in these contexts may provide a more holistic view of how tissue and CSF dynamics coordinate to promote brain clearance.

## Acknowledgements

We would like to thank Estee Perelgut, Sarah Richter, Kyle Droppa and Julianna Gerold for their help with subject recruitment and MRI scanning support, Azma Mareyam for the use of her inhouse-built 7T RF coil, Jason Stockmann for his help with the phantom construction, and Prof. Simon Robinson for use of the ASPIRE pulse sequence and image reconstruction “C2P” package. This work was supported in part by the NIH NIBIB (grants P41-EB030006, R01-EB019437, and R01-EB036507), NCCIH (grant R01-AT011429), NINDS (grant F31-NS141336), NIA (grant R00-AG083056), by the *BRAIN Initiative* (NIH NINDS grants U19-NS123717, U19-NS128613, and U24-NS129893), and by the MGH/HST Athinoula A. Martinos Center for Biomedical Imaging; and was made possible by the resources provided by NIH Shared Instrumentation Grant S10-OD023637.

---

**Figure S1.**
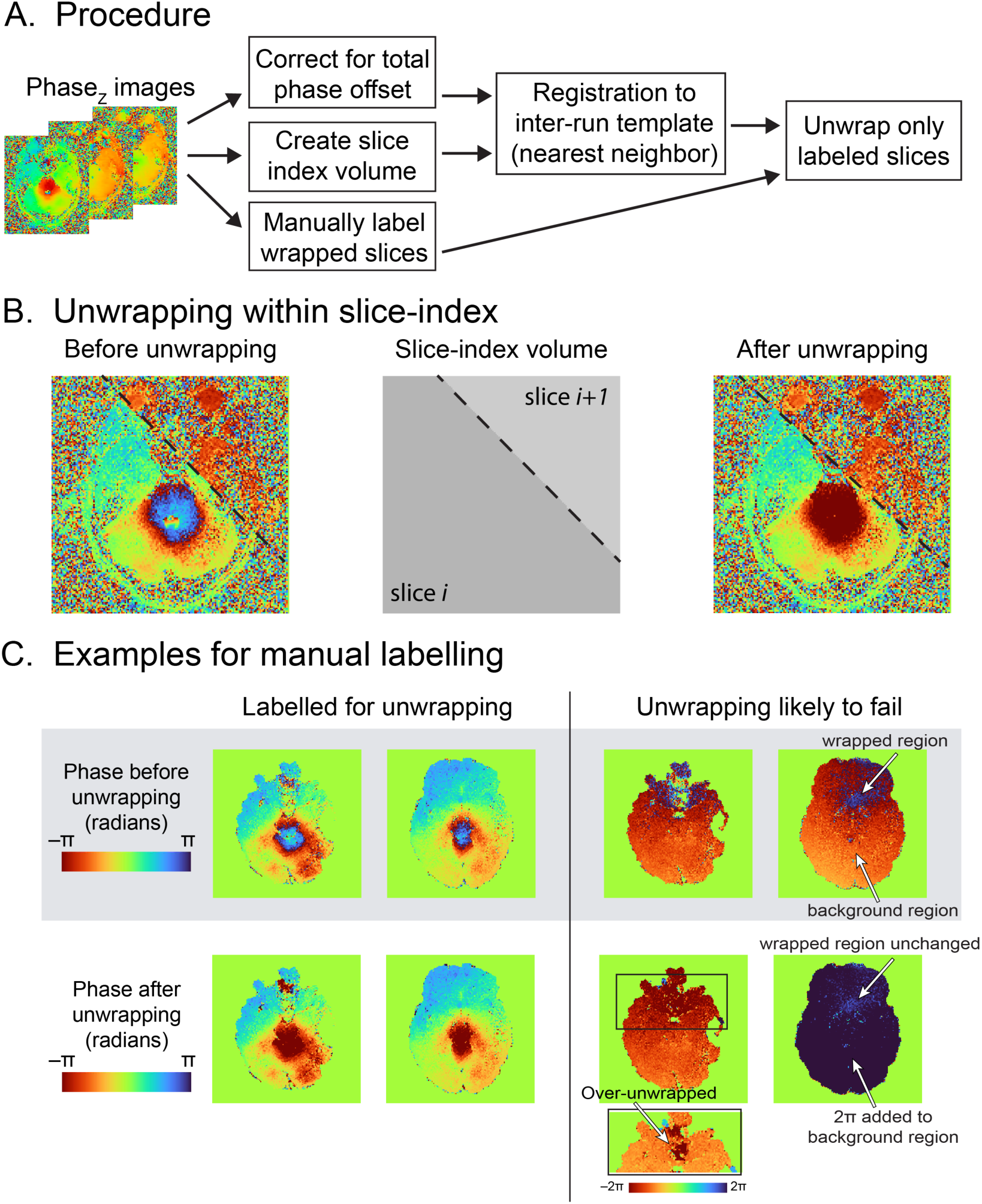
Processing and manual labeling procedure for phase unwrapping. (A) Summary of pipeline for unwrapping within acquired slices, including manual labeling. (B) Example slice after registration and before unwrapping, where the slice boundary intersects the image. Within slice-index unwrapping avoids treating the slice boundary as a phase wrap. (C) Comparison of individual slices that were manually selected (left) or not selected (right) for phase unwrapping using example data from the 2×2×2 mm^3^ protocol. The slices labelled for unwrapping have centralized, focal areas of wrapped phase and are correctly unwrapped by the 2D algorithm. The slices that were not selected exhibited noisy, non-centralized areas of wrapped phase and result in “over-unwrapping” or are otherwise poorly unwrapped.

**Figure S2.**
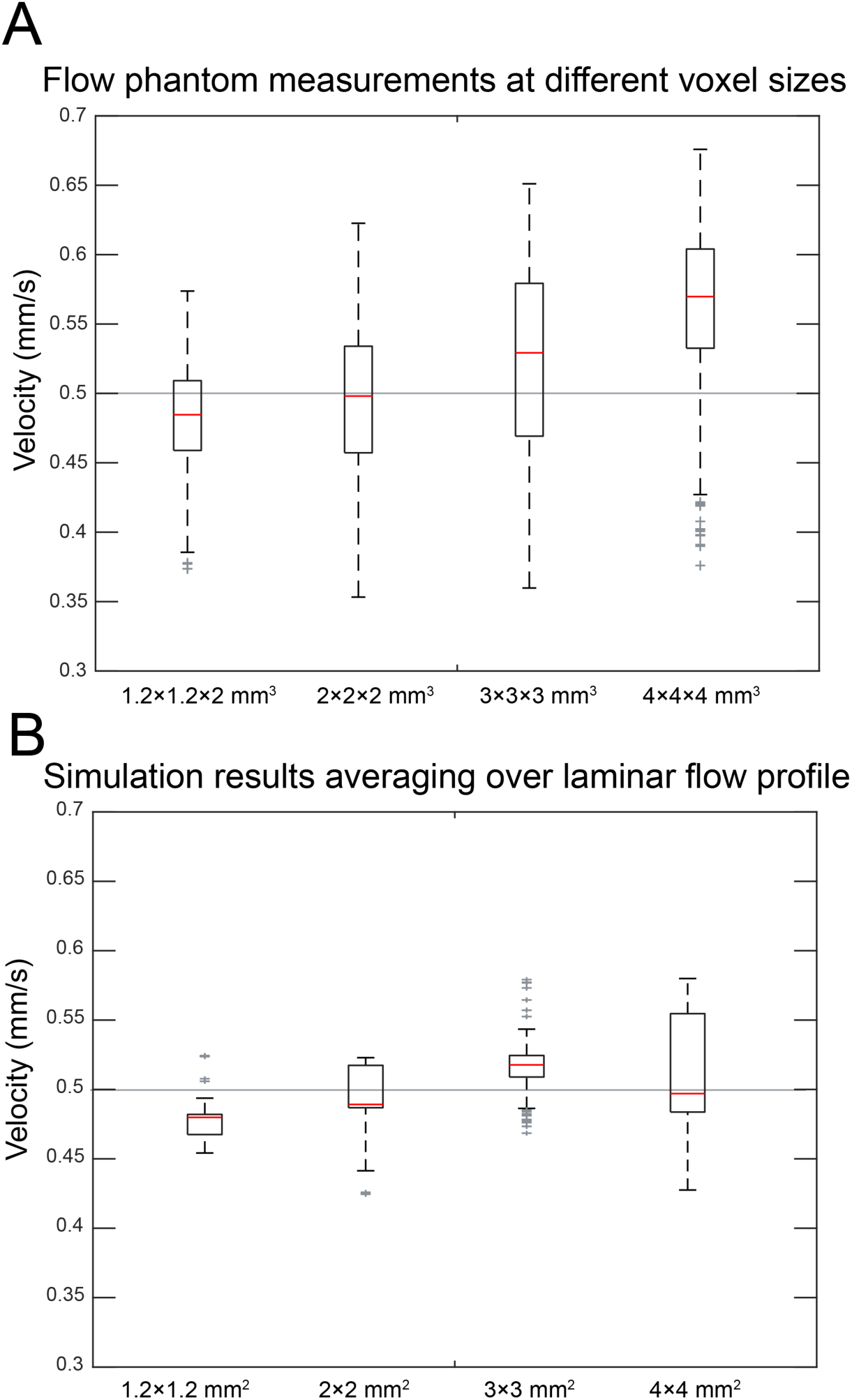
Velocity estimation biases in our flow phantom may be due to effects of a laminar flow profile. (A) Flow phantom velocity estimates using the four protocols corresponding to the four voxel sizes used for *in vivo* measurements. Each sample is the velocity estimate from a single tube (inner diameter = 6.4 mm). True mean velocity is 0.5 mm/s. A voxel-size-dependent bias is seen, which results in underestimation for smaller voxels and overestimation for larger voxels. (B) Simulations of the effects of 2D spatial averaging over a laminar flow profile with matched tube diameter and mean velocity reproduces the voxel-size-dependent velocity estimation bias. The simulation was conducted by systematically shuffling the position of the voxel grid over the tube cross-section to account for different spatial sampling of the tube in each slice. Each simulated velocity estimate corresponds is the average velocity from one such position. Note the similarity in the average measured velocity between phantom measurements and simulations. The variability in velocity estimates is highest in the largest voxel size for both flow phantom measurements and laminar profile simulations. Average velocity in voxels that are small relative to the laminar flow profile will be biased towards the slower velocities at the tube interface (caused by friction) resulting in underestimated velocity; voxels that are large compared with the laminar profile will be biased by the peak velocity resulting in overestimation.

**Figure S3.**
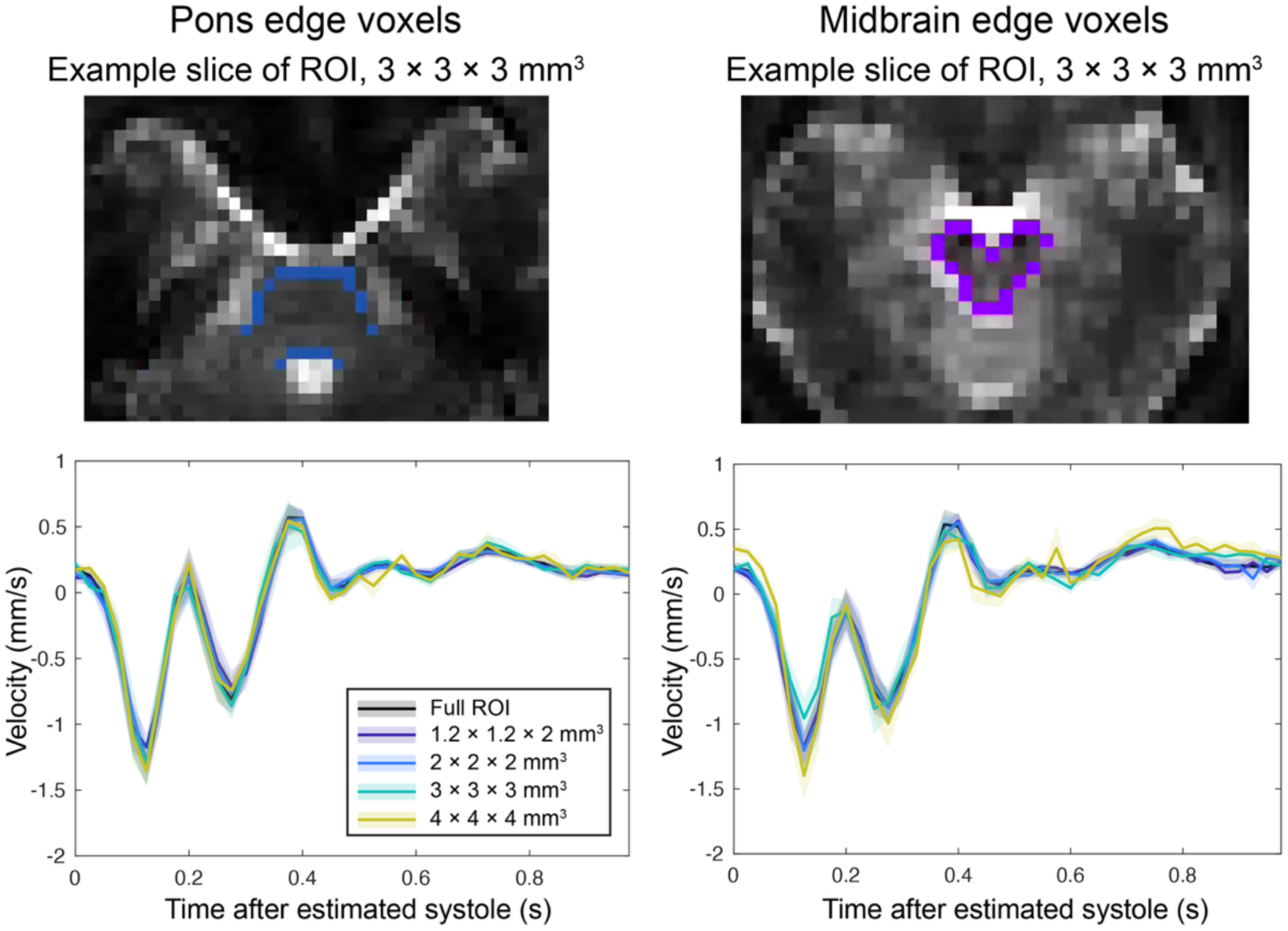
Cardiac-locked velocity patterns in voxels with high tissue-CSF partial volume effects. Pons and midbrain ROIs were modified to only include voxels at the edge of the structure, adjacent to CSF; see images for example “edge-voxel” ROIs. Cardiac-locked velocity time courses from edge-voxel ROIs are plotted for each voxel size to assess partial volume-induced bias in velocity estimations, with the average response in the full ROI included in black for comparison. Error bars indicate standard error across subjects.

